# Why are viral genomes so fragile? The bottleneck hypothesis

**DOI:** 10.1101/2021.01.23.427881

**Authors:** Nono S. C. Merleau, Sophie Pénisson, Philip J. Gerrish, Santiago F. Elena, Matteo Smerlak

## Abstract

If they undergo new mutations at each replication cycle, why are RNA viral genomes so fragile, with most mutations being either strongly deleterious or lethal? Here we provide theoretical evidence for the hypothesis that genetic fragility evolves as a consequence of the pervasive population bottlenecks experienced by viral populations at various stages of their life cycles. Modelling within-host viral populations as multi-type branching processes, we show that mutational fragility lowers the rate at which Muller’s ratchet clicks and increases the survival probability through multiple bottlenecks. In the context of a susceptible-exposed-infectious-recovered epidemiological model, we find that the attack rate of fragile viral strains can exceed that of more robust strains, particularly at low infectivities and high mutation rates. Our findings highlight the importance of demographic events such as transmission bottlenecks in shaping the genetic architecture of viral pathogens.

## Introduction

From tobacco mosaic virus to poliovirus and SARS-CoV-2, some of the most consequential plant, animal and human pathogens are RNA viruses. In spite of their extremely small genomes, these organisms find adaptive solutions to environmental challenges such as hosts’ immune response, pervasive differences in susceptible cell types, switches in host and vector species, and antiviral drugs (1–3). The remarkable evolvability of RNA viruses has been linked to their error-prone replication, short generation times, and large population sizes (4). But high mutation rates are a double-edged sword: while replication errors provide the fuel necessary for rapid adaptation, they also increase the genetic load on viral populations, which in turn imposes a limit to genome size (5, 6). Moreover, evidence gathered from diverse viral systems (7–11) shows that high mutation rates coupled with strong population bottlenecks (*e.g*. associated with airborne or fomite transmission events) turn on Muller’s ratchet (12), resulting in the loss of fit genotypes (13, 14). As fitness declines, populations risk experiencing a mutational meltdown, with low fitness genotypes unable to restore large population sizes and deleterious mutations accumulating at an ever increasing rate (15, 16). How do RNA viruses manage to persist in the face of these challenges?

RNA viruses may have evolved specific mechanisms to maintain genome integrity in the face of high mutation rates (17–20). Proposed mechanisms include complementation at high multiplicity of infection during transmission *e.g*., by physically aggregating viral particles (21, 22); the use of stamping machine, rather than geometric, replication mechanisms (23); the segmenting of viral genomes with segment reassortments during mixed infections or increased recombination rates, two simple forms of sex that reduce mutational load (24, 25); or the co-opting of cellular chaperones (*e.g*., heat-shock proteins) to assist at different stages of the replication cycle (26). It is possible that all of these mechanisms (and others yet to be discovered) play a role in mitigating the damage done by mutations. Indeed, general arguments suggest that neutral evolution tends to reinforce mutational robustness by moving viral populations away from the edges of neutral networks (27–29).

Somewhat paradoxically, the opposite strategy of *maximizing* mutational damage may also be a key component of the evolutionary response to low-fidelity replication. Both empirical and theoretical arguments support this hypothesis. Firstly, viral RNA genomes generally contain overlapping reading frames, encode for polyproteins that need to be precisely processed post-translationally, and express multifunctional proteins involved in different processes along the infection cycle; these structural properties predict large deleterious effects of most mutations (5, 6), as is indeed observed (30). Secondly, analyses of Muller’s ratchet show that the risk of mutational meltdown is highest when deleterious effects are moderate (16). This is because weak deleterious mutations negatively impact population fitness without being strongly selected against (Fig. 1). This observation has led several authors to the conclusion that genetic fragility is in fact selected for in the high mutation rate regime of evolution (31–33).

**Fig. 1.**
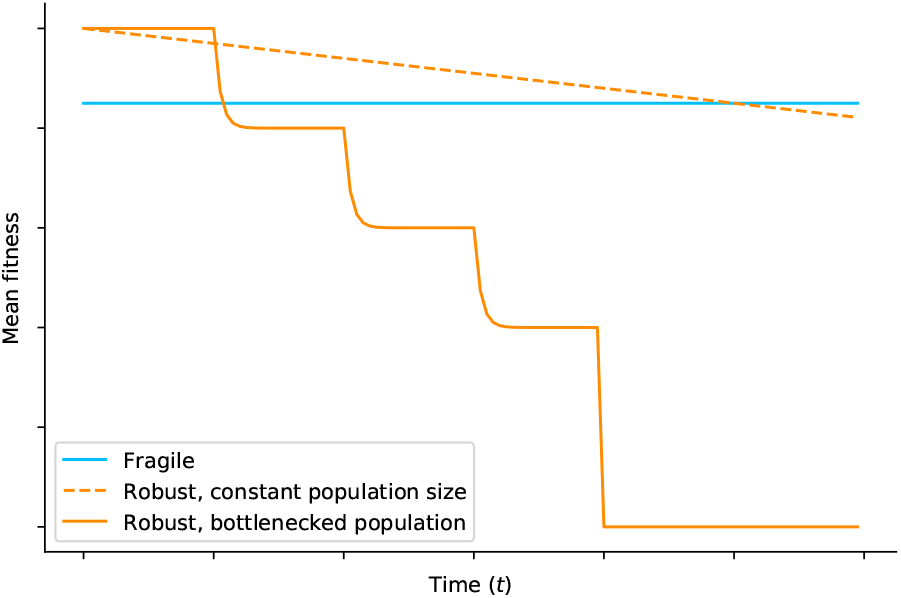
Schematic of fitness dynamics. Robust genomes (orange) initially have a lower mutational load, and therefore higher population fitness, than fragile genomes (blue). But they also fix deleterious mutations more frequently, leading to a gradual decline in population fitness (dashed line); this effect is more pronounced when bottlenecks weaken the strength of selection and accelerate the clicking of Muller’s ratchet (continuous line).

In this paper we show that the genetic fragility of RNA viruses may be explained by the population bottlenecks they experience during their cycle. Our contribution is twofold. First, we model population bottlenecks (and the genetic drift they induce, including Muller’s ratchet) explicitly using multi-type branching processes (34, 35); such bottlenecks are not easily modelled within more common approaches based on weak-mutation strong-selection limit (32) or quasi-species theory (31, 33). Using general results in branching process theory, we derive expressions for the survival probability of a population through repeated bottlenecks as a function of the deleterious effect of mutations, confirming that fragility can be advantageous in the long run. Second, we consider the epidemiology of genetic fragility using a simple agent-based compartmental model. Evolutionary epidemiology (36) is an emerging field focusing on the interactions between evolutionary and epidemiological dynamics that seeks to explain the evolution of virulence and other properties of pathogens. Here we ask under what epidemiological conditions a fragile strain can have a higher attack rate than a robust one. We find that two parameters determine whether or not this is possible: the mutation rate *u* and the infection rate *β* (both relative to the recovery rate), with high *u* and low *β* both favouring fragile viruses.

Our model builds upon the standard model of Muller’s ratchet (13), under which all mutations have the same deleterious effect (in particular, none is lethal), and do not interact epistatically. Positive epistasis among deleterious mutations has in fact been shown to be a pervasive phenomenon in compacted RNA genomes (37). Furthermore, the fraction of mutations that are lethal is usually large for RNA viruses (38). We rationalize these simplifications by noting that relaxing them would further increase the long-term advantage of fragile genomes, reinforcing the argument for the bottleneck hypothesis.

## Results

### Muller’s ratchet in expanding viral populations

Consider a small viral population with absolute fitness *w*_0_ > 1 and genomic mutation rate *u*. Assume that all mutations are deleterious with the same effect *s_d_*, such that an individual carrying *i* mutations—an “*i*-mutant”—has fitness *w_i_* ≡ *w*_0_ (1 − *s_d_*)^*i*^. The dynamics of such a population falls under two broad alternatives: either it goes extinct through demographic fluctuations, or it grows to unbounded sizes with an asymptotic mean fitness 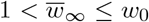; the latter outcome is only possible if *w*_0_*e*^−*u*^ > 1. This is illustrated in Fig. 2 for a low value of *s_d_* (a “robust” type) and a high value of *s_d_* (a “fragile” type).

**Fig. 2.**
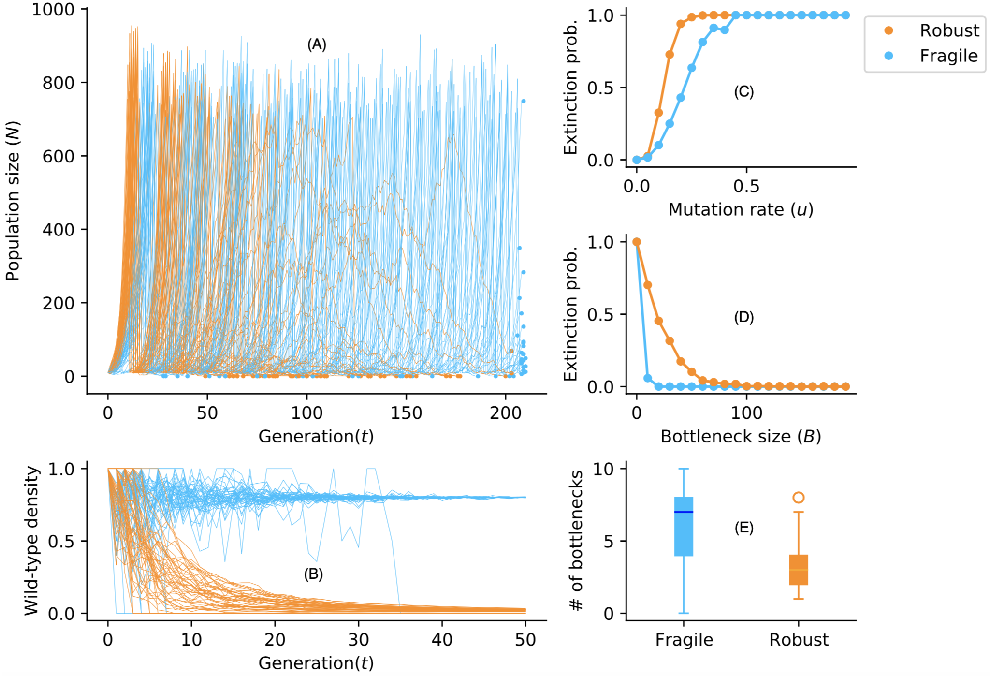
Viral populations through multiple bottlenecks. For an initial population size *N* = 10 with fitness *w*_0_ = 1.5 and a mutation rate *u* = 0.2, viral particles reproduce, mutate, and die out. Once the populations reach the carrying capacity *C* = 800, a sub-population of size *B* = 10 is sampled and the branching process is restarted. (*A*) Populations of robust viruses grow faster but go extinct more often after multiple bottlenecks. (*B*) The wild-type (individual with fitness *w*_0_) density in the populations over time. (*C*) Effect of mutation rate on the extinction probability for a fixed bottleneck size *B* = 10. (*D*) Effect of *B* for a fixed mutation rate (*u* = 0.2) on the extinction probability. (*E*) Number of bottlenecks before extinction.

Key to the fate of the population is the onset of Muller’s ratchet, *i.e*. the extinction of fit genotypes through genetic drift in small populations. The ratchet mostly clicks during the early states of the expansion, when the population size is smallest and extinction is likely. But it may also start clicking later through some rare fluctuation; if this click is followed by another click, and then another, the population can start shrinking again into mutational meltdown. If the ratchet clicks too many times, namely more than *K* = max{*k*: *w_k_e*^−*u*^ > 1} times, extinction is unavoidable: any mutant carrying more than *K* mutations has absolute fitness smaller than one, and therefore generates a subcritical branching process.

Robust and fragile populations experience Muller’s ratchet differently. For a robust genotype, mutations have a small deleterious effect, and are therefore under weak negative selection. This makes a click of the ratchet quite likely. On the other hand, each click comes at a low fitness cost, and so the population can withstand a relatively large number of clicks. Fragile populations, by contrast, are under strong negative selection against deleterious mutations, hence clicks are rarer; when they do occur, though, extinction becomes almost certain. Which fares better?

Using branching process theory we compute analytically the probability *p_k_* that the fittest surviving individual carries exactly *k* mutations, *i.e*. that Muller’s ratchet click *k* times (Eq. (1) in Materials and Methods). From this, we find that both the survival probability 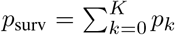 and the expected asymptotic population mean fitness 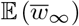 (Eq. (2)) are decreasing functions of *s_d_*, consistent with the idea that, when all mutations are deleterious, mutational robustness is an evolutionary advantage. But there is a caveat: because Muller’s ratchet clicks less frequently when genomes are fragile, the populations that do emerge from the expansion tend to be mutations-free. As a result, the asymptotic mean population fitness *conditional on survival* 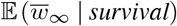 turns out to be non-monotonic in *s_d_*, and in fact to be maxi-mized for large deleterious effects (Fig. 3).

**Fig. 3.**
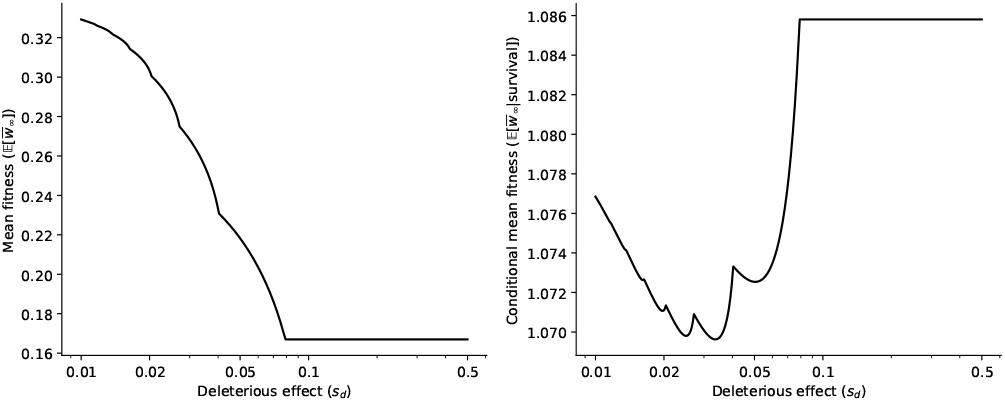
Analytical results for expected asymptotic population mean fitness, both unconditional (left) and conditional on survival (right) as a function of the deleterious effect of mutations *s_d_*, for a population with *w*_0_ = 1.2 and *u* = 0.1. While a more robust strain has a lower mutational load and therefore a higher population mean fitness (left), it is also more vulnerable to Muller’s ratchet, implying that surviving populations tend to have lower mean fitness (right).

### Survival of viral population through multiple bottlenecks

The adaptive value of virus’ mutational fragility becomes apparent when we consider a succession of bottleneck-expansion cycles (Fig. 2), corresponding to viral transmission followed by within-host replication. This process can be modeled as a Markov chain on the space of post-bottleneck populations. Let *B* denote the size of the population after the transmission bottleneck. (In some cases, *B* can be a small as 1 (39, 40)).

We model virus transmission as sampling without replacement from a surviving population as described above, resulting in a new founding population with composition **n** = (*n*_0_,⋯, *n_K_*), where *n_i_* is the number of *i*-mutants and we omit the subcritical mutants whose lineage will go extinct with probability one (hence 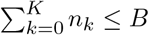). After within-host expansion and transmission sampling, this population will give rise to a new founding population with composition **m** = (*m*_0_,⋯, *m_K_*) with a probability *P*(**n** → **m**) given explicitly in Eq. (15) in Materials and Methods. From this Markov chain, we can compute the probability that a population can survive any given number of bottlenecks (Eq. (4)). Fig. 4 shows the survival probability after up to five bottlenecks of size *B* = 5 as a function of the deleterious effect *s_d_*. Although non-monotonic, this probability is maximized at large *s_d_* when the number of bottlenecks increases, *i.e*. extinction becomes less likely for more fragile genomes. The results in the previous paragraph explain why: fragile populations are protected against Muller’s ratchet, hence each new infection starts from fit founders. By contrast, robust genomes accumulate deleterious mutations; after several transmission bottlenecks, the founder particles tend to have low fitness and become increasingly unlikely to give rise to surviving lineages. It is as if robustness promoted mutational meltdown on a longer time scale—a meta-population meltdown.

**Fig. 4.**
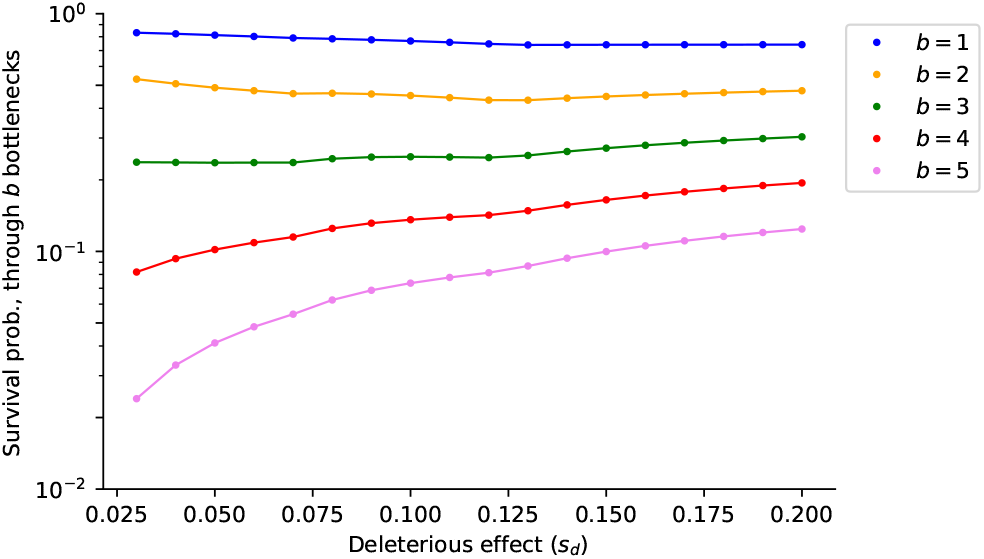
Analytical results for the survival probability *vs*. mutational fragility for a population with *w*_0_ = 1.2 and *u* = 0.05 undergoing *b* bottlenecks of size *B* = 5, computed using the bottleneck-to-bottleneck Markov chain *P*(**n** → **m**). The higher the number of bottlenecks a populations has to survive, the more fragility is selected for.

### Epidemiology of fragility

How would these effects play out in an epidemic outbreak? Would the lower propensity of fragile genomes to suffer meltdown make up for their lower initial population fitness? To investigate this question we consider a Susceptible-Exposed-Infectious-Recovered (SEIR) model defined as follows. When a susceptible *S* meets an infectious individual *I*, a sample of the latter’s viral population with size *B* is transmitted and *S* becomes exposed *E*. After this event, the within-host viral population carried by *E* can either (*i*) go extinct, in which case *E* returns to the susceptible compartment (*E* → *S*) or (*ii*) grow exponentially during an incubation period *τ* until it reaches a critical threshold *C* which makes the host infectious (*E* → *I*). Which is more likely depends on the viral genetic parameters (*w*_0_, *s_d_*, *u*) and, as shown in the previous section, on the numbers of supercritical viral particles transmitted to the new host, described by some vector **n**. The incubation time is random as well, with a probability distribution depending on **n** and on the number of clicks of Muller’s ratchet during the expansion, see Eq. (5). (We show in *SI text* that this distribution can be approximated by a Gamma distribution depending on these parameters.) The host population is assumed to be well-mixed (no spatial structure): at each time step and for each pair (*S, I*), transmission occurs with a probability *β*. Recovery in turn takes place at a rate *γ*.

Fig. 5 presents the results of simulations where a susceptible population of size *N* = 5000 is seeded with 100 infectious individuals, half of which carry a robust viral strain (*s_d_* = 0.05) and the other half a fragile one (*s_d_* = 0.9). Three regimes emerge depending on the transmission rate *β* and the mutation rate *u* (at fixed bottleneck size *B* = 5). When infectivity is high and the population quickly becomes completely infected, the robust strain fares better due to its higher initial fitness and faster growth rate. When infectivity is low, however, fragile strains prove to have a much higher attack rate. (We define the attack rate of a strain as the fraction of the host population which is exposed, infectious or recovered with that strain.). As noted in Fig. 5*A*, the robust strains are more virulent in the early stages of the epidemic. But as time goes on, the robust genomes loose fitness and increase their post-bottleneck extinction probabilities. That is also the reason why there is a small decrease in the total number of robust genome infections in (*E* + *R* + *I*)-populations, as exposed hosts return to the susceptible compartment (Fig. 5*A*). At a late stage (*t* > 200), mutationally robust viruses are too degraded to make their host infectious. Finally, very high mutation rates and low infectivities do not allow for the epidemic to spread, as all viral populations (robust and fragile) quickly go extinct. These findings are summarized in Fig. 5*D* in terms of the median ratio of attack rates across 50 runs for each parameter set.

**Fig. 5.**
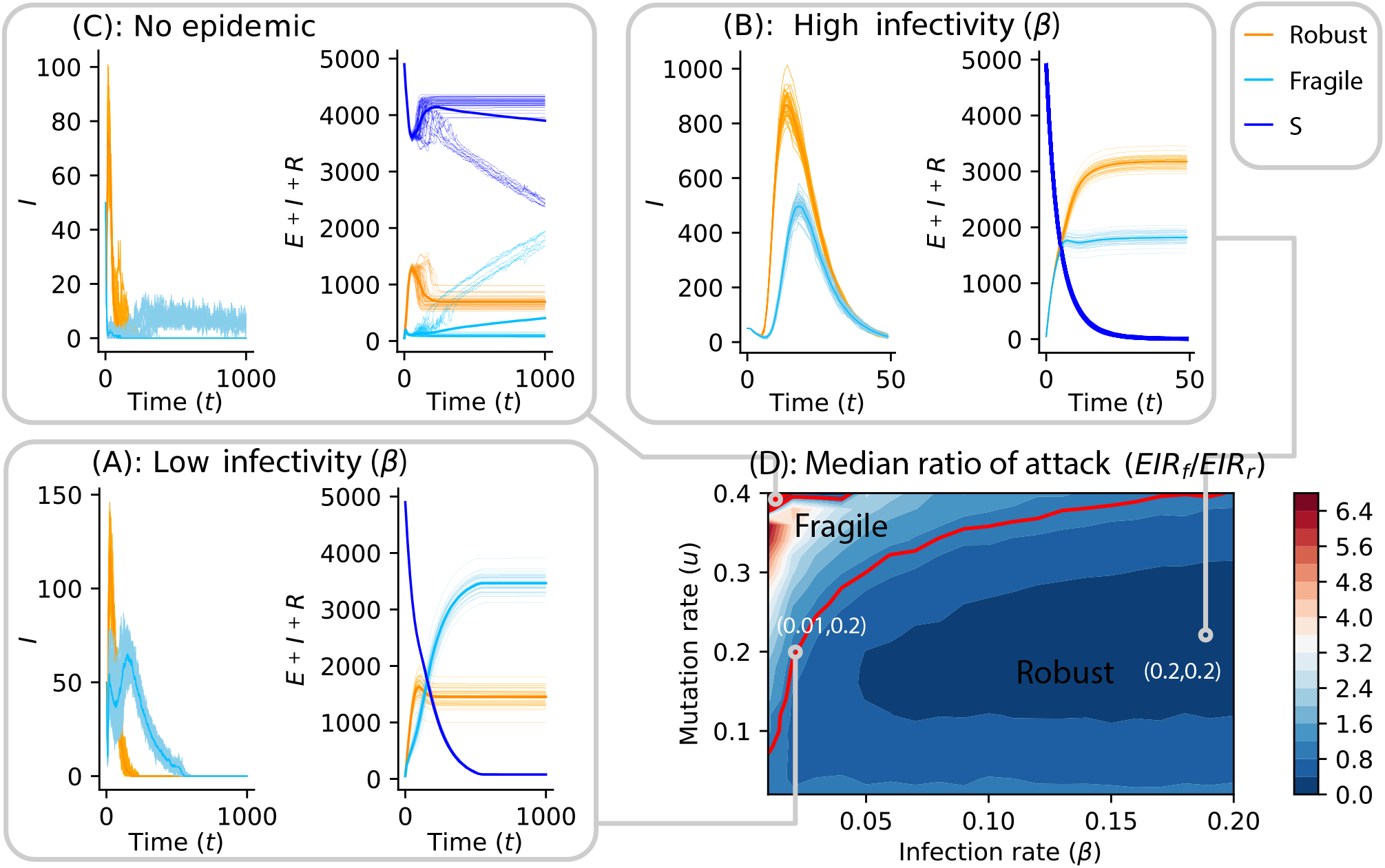
Agent-based simulation of the SEIR model. (*A*) At low infection rate (*β* = 0.01) and high mutation rate *u* = 0.2, the robust genomes are more infectious in the beginning of the epidemic than the fragile ones, but as time goes on, it turns out that the fragile genomes become more virulent. (*B*) Using the same mutation rate and the highest infectivity gives the robust genome a chance to become the most infectious because it reproduces faster, and its population undergoes fewer bottlenecks. (*C*) At low infection rate and very high mutation rate (*u* = 0.4), the disease does not spread in the population, due to the high extinction probability ofthe viral populations. (*D*) Median ratio of attack rate (which is the total number of fragile infections divided by the robust ones) across 50 runs for each (*β, u*) parameters.

Underlying this contrast is the distribution of incubation times, which differs for the robust and fragile strains (Fig. 6*A* and Fig. 6*B*). The same observation can be drawn by looking at the mean basic reproduction number of both robust and fragile strains at low infectivity (and respectively at high infectivity) (see Fig. 6*C* and Fig. 6*D*). Over time, the basic reproduction number *R_t_* of fragile strains stays constant while the robust one decreases.

**Fig. 6.**
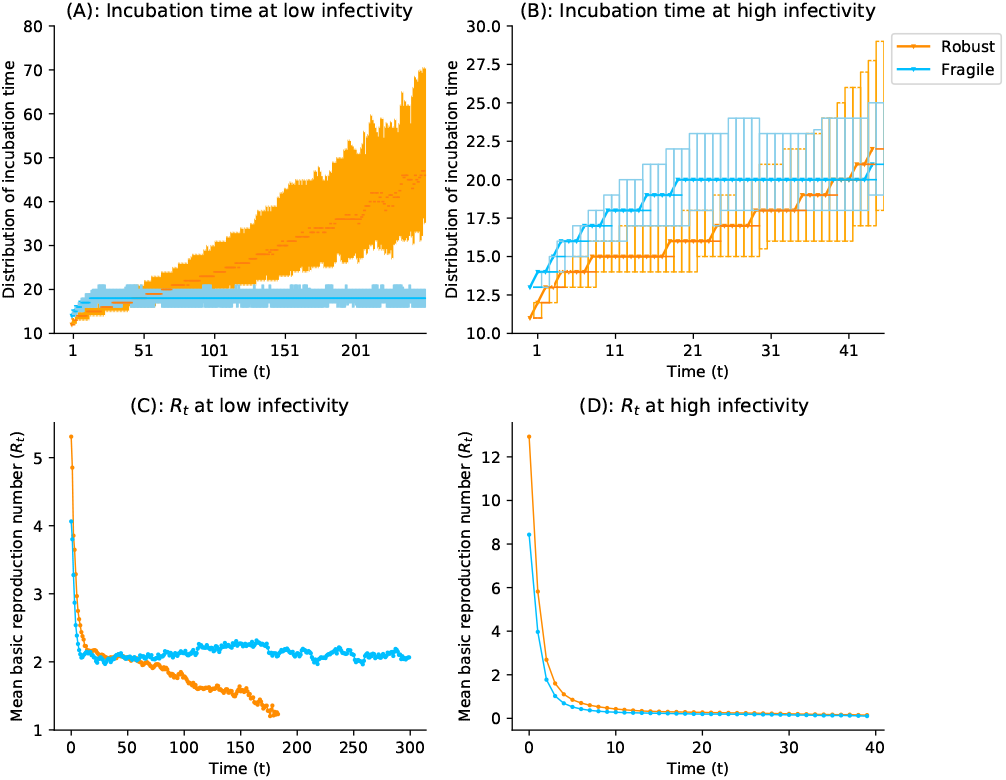
Incubation time distributions and mean basic reproduction number at low infectivity (*β* = 0.01) (respectively at high infectivity (*β* = 0.2)), with viral parameters (*w*_0_ = 1.5, *u* = 0.1) and bottleneck size *B* = 5. (A) The epidemic lasts longer and the robust viral populations lose fitness. The later transmitted viral particles take a longer time to grow up to *C*. In contrast, the incubation times of the fragile strains stay on average constant. (*B*) On short-time epidemic, the mean incubation time of the robust strain stays smaller than the fragile ones. (*C*) The epidemic lasts longer, wherein the beginning, the reproduction number of the robust strain is higher than the fragile one but later on, when the robust strain’s Rt decreases, the fragile one stays constant. (*D*) The virus takes over the population in the early stage of the epidemic when the robust strains are the most virulent.

## Discussion

There is a growing appreciation for the fact that population bottlenecks are not just a fundamental aspect of the life cycle of viruses, but that they also play a key role in their evolution (41, 42). The effect of bottlenecks is not always detrimental: bottlenecks can effectively remove cheaters, *e.g*. defective interfering viruses (43), or enhance the effectiveness of selection if beneficial alleles act in *trans* (44). In addition, genetic bottlenecks can facilitate traveling across the characteristically rugged fitness landscapes of RNA viruses (37), where it is easy for viruses to become trapped at suboptimal fitness peaks (45, 46); by relaxing the intensity of selection, bottlenecks enable the exploration of new regions of the landscape.

In this paper we have explored another aspect of highly mutable populations subjected to periodic bottlenecks: they experience a strong evolutionary pressure towards genetic fragility. Earlier work has established that intermediate deleterious effects *s_d_* maximize the strength and speed of Muller’s ratchet (16, 31); similar results have been reported more recently in terms of “ratchet robustness” (47) or “drift robustness” (32). We find this U-shaped pattern in the context of expanding populations as well, *e.g*. for the asymptotic population mean fitness given survival (Fig. 3); the unconditional population mean fitness, by contrast, always decreases with fragility, which is consistent with another analysis of the advantage of mutational robustness (33). By modelling changes in population sizes with branching processes, we allowed a more complete picture to emerge. In this picture, the evolution of high neutrality (27, 28) is not incompatible with selection for maximally deleterious mutations (31) and anti-redundancy (33). An agent-based SEIR epidemiological model further reveals the determinants of genetic fragility, highlighting the importance of epidemic transmission parameters.

The evolution of viruses is often described in terms of fitness landscapes and their topographies. Our findings highlight the limitation of this picture: the motion of evolving population in genotype space depends on the structure of the genotype-phenotype-fitness mapping, but also on mutation rates and life cycle parameters such as the frequency and stringency of population bottlenecks. As a result, the evolution of mutational robustness—or mutational fragility—cannot be construed solely as the search for an optimal region in the fitness landscape, be it the highest peak or the flattest plateau. Our multi-type branching process approach is suggestive that a composite definition of fitness might be more predictive of evolutionary success in the present context, namely, a definition that takes account of both offspring number (Malthusian fitness) and long-term survival probability. The augmented evolutionary relevance of this definition to the present context is manifest in the comparison between the left and right panels of Fig. 3.

To be sure, our model relies on simplifying assumptions, mainly pertaining to the nature of underlying mutational landscapes and to epidemiological details. The assumption that all mutations having the same deleterious effect is a common one that simplifies the mathematics. Fitness effects of deleterious mutations are more realistically modeled as a random variable with a continuous, heavy-tailed distribution (*e.g*. the Gamma and Weibull distributions have been previously used to satisfactorily fit experimental data (48, 49)) or a U-shaped distribution to incorporate lethal mutations. As already noted, we make another common assumption that does not necessarily hold for real viral genomes, namely, the independence of mutational effects. Evidence pervasively suggests that positive epistasis is the norm for compacted viral RNA genomes (37), including many instances of compensatory mutations. Indeed, it was shown long ago that if deleterious alleles interact synergistically, they are more efficiently removed from the population and thereby slow the advance of Muller’s ratchet (50). Finally, our evolutionary epidemiology approach ignores co- or super-infection cases wherein multiple viral strains infect the same host. On the one hand, multiple infections allow competition among strains to occur at the level of individual lineages. On the other hand, multiple infections allow the sharing of gene products among different genotypes within a cell, thus compensating for deleterious effects and eventually contributing to the accumulation of more mildly deleterious mutations in the population (51). In its current form, our model may well serve as the null against which to test the effect of all these important factors in the fate of evolving viral populations.

Overall, our findings can be summarized with a classic metaphor. Robust genomes are hares: they grow fast but accumulate mutational damage through Muller’s ratchet, which jeopardizes their potential for long-term survival. Fragile genomes are turtles: they grow more slowly but weather bottlenecks more reliably and have higher long-term survival probabilities. Turtles may seem weak individually, but as a group they have survived for hundreds of millions of years. Similarly, fragile viral genomes are individually vulnerable to deleterious mutations; at the meta-population level, however, they may hold the key to evolutionary resilience.

## Materials and Methods

The evolution of viral populations experiencing Muller’s ratchet and going through bottlenecks is modeled by a multitype branching process, in which a “type” corresponds to the number of accumulated deleterious mutations. We refer to *SI text* for a detailed description of the model and a rigorous demonstration of the following results.

Starting with **n** supercritical mutants, the extinction probability *p*_ext,*k*_(**n**) of all types up to type *k* in the population is given by the element-wise product 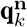, where **q**_*k*_ is the smallest solution to a fixed-point equation involving the generating function of the branching process. Muller’s ratchet click probability *p_k_*(**n**) that the fittest surviving individuals carry *k* mutations is then

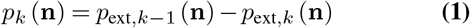

and the population survival probability 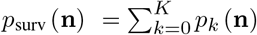.

On the event that the fittest surviving individuals carry *k* mutations, the asymptotic mutant spectrum of the population is given by (0,…, 0, *f*_0_, *f*_1_,…), where *f_i_* is the Poisson frequency *f_i_* = *e*^−*u*/*s_d_*^(*u/s_d_*)^*i*^/*i*!. The asymptotic mean population fitness is thus a random variable 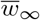 equal to *w_k_e*^−*u*^ with probability *p_k_*(**n**) for 0 ≤ *k* ≤ *K*, and to 0 otherwise.

Therefore

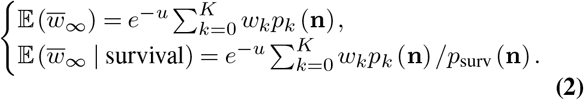

When going through a bottleneck, this population gives rise to a new founding population with composition **m**, according to transition probability

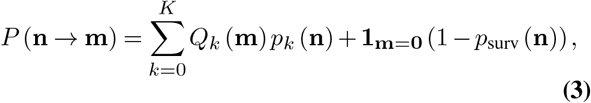

where *Q_k_*(**m**) is the probability of getting **m** supercritical mutants in the sample of size *B*. The resulting transition matrix *P* then provides an explicit expression of the probability for a viral population with initial state **n** to become extinct after going through at most *b* bottlenecks:

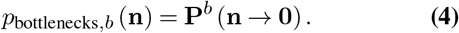

In our SEIR epidemiological model, the time spent in the exposed state depends on **n** and on the number of Muller’s ratchet clicks during the expansion, or equivalently on the type *k* of the fittest surviving individuals. We show that this incubation period *τ*_**n**,*k*_ is

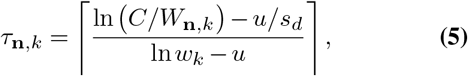

where *W*_**n**,*k*_ is a positive random variable and ⌈·⌉ stands for the ceiling function. As exposed in *SI text, W*_**n**, *k*_ can be expressed as the sum of a random number of independent exponential random variables with common parameter

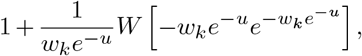

where *W* stands for the principal solution of the Lambert function.

Simulations were carried out using an agent-based approach in Python. We have created a repository at github.com/strevol-mpi-mis/EvoEpi containing the code and minimum documentation.

## ACKNOWLEDGEMENTS

We thank the members of the Structure of Evolution group at MPI MiS for useful discussions. Funding for this work was provided by the Alexander von Humboldt Foundation in the framework of the Sofja Kovalevskaja Award endowed by the German Federal Ministry of Education and Research to M.S. Work in València was supported by Spain Agencia Estatal de Investigación - FEDER grant PID2019-103998GB-I00 and Generalitat Valenciana grant PROMETEO2019/012 toS.FE.

## Supplementary Note 1: Muller’s ratchet in expanding populations

### Branching process setting

The composition of the viral population is determined by the number of virions carrying 0, 1, 2, *etc*. deleterious mutations. Its evolution is modeled by a branching process (52): each viral particle in generation *n* randomly produces particles in generation *n* + 1, independently of the rest of the population and according to some probability distribution which depends solely on the “type” of the particle. Since there are no back mutations, the resulting process 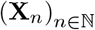 is a reducible branching process with infinitely many types, where 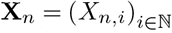 is the population composition at time *n*, and *X_n,i_* the number of *i*-mutants (particles carrying *i* mutations) at this time. All the properties of the branching process are contained in its offspring generating functions *F*_0_, *F*_1_,…. The generating function *F_i_* determines the distribution of the number of various mutants to be produced by an *i*-mutant. By definition, for 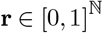, *F_i_*(**r**) is the expected value of 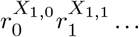 when the process is initiated by one *i*-mutant. Because there are no back mutations, it solely depends on *r*_*i+j*_, *j* ≥ 0, and we write *F_i_*(*r_i_, r*_*i*+1_,…).

We assume throughout this paper that the average number of (*i* + *j*)-mutants produced by an *i*-mutant is *w_i_e*^−*u*^*u^j^*/*j*!. This is achieved for instance if the total number of offspring of an *i*-mutant follows a Poisson distribution 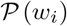, and if each of its offspring accumulates a number of additional deleterious mutations according to 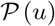. In this particular case, the *i*-th generating function is

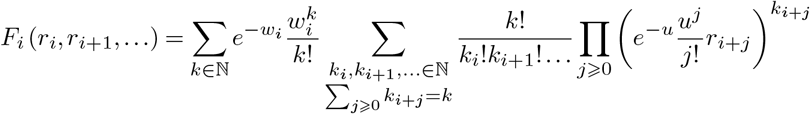

which simplifies to

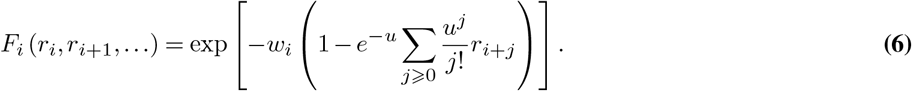

### Survival probability

Let *m_i_* = *w_i_e^−u^* the average number of *i*-mutants produced by an *i*-mutant. It follows from a general result of branching processes theory that if *m_i_* ≤ 1, any population starting with one *i*-mutant is doomed to extinction; if *m_i_* > 1, this population has positive survival probability. Let *K* = max{*k*: *m_k_* > 1} the largest number of mutations which can be carried by an individual without leading to its progeny’s extinction, and call *supercritical* any *k*-mutant with *k* ≤ *K*. By independence of the lineages, the fate of the population 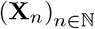 only depends on the initial numbers of supercritical mutants **n** = (*X*_0,0_,…, *X*_0,*K*_). First, we compute for each *k* ≤ *K* the *partial* extinction probability *p*_ext,*k*_(**n**) of all mutants up to type *k*, given the initial condition **n**, namely the probability of {*X_n,i_* → 0, ∀*i* ≤ *k*}. It corresponds to the extinction probability of the reducible branching process with finite set of types {0, 1,…, *k*} and generating functions *F_i_*(*r_i_*,…, *r_k_*, 1,1,…), *i* ≤ *k*. As such (53), the partial extinction probability is given by

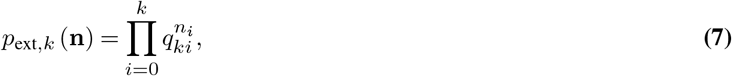

where (*q*_*k*0_,…, *q_kk_*) is the smallest nonnegative solution of the system of equations

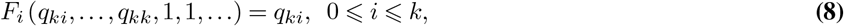

(see Eq. (24) for an explicit solution when *F_i_* is given by Eq. (6)). The overall survival and extinction probabilities of the population are then obtained as

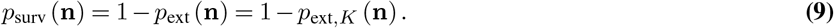

### Muller’s ratchet click probabilities

For a population starting with supercritical mutants **n** and *k* ⩽ *K*, let *p_k_*(**n**) the probability that the fittest surviving individuals carry *k* mutations. Equivalently, it is the probability that Muller’s ratchet clicks *k* – *k*_0_ times, where *k*_0_ = min{*i*: *n_i_* > 0} is the fittest initial supercritical type. It corresponds to event 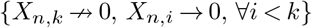 (all *i*-mutants with *i* < *k* go extinct but *k*-mutants survive), therefore

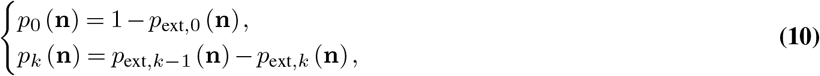

It comes in particular 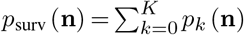, consistent with the fact that any click beyond the threshold *K* leads to extinction.

### Mutant spectra and asymptotic mean fitness

The late-time composition of the population is completely determined by the number of clicks of the ratchet. On the event Ω_*k*_ that the type of its fittest surviving individuals is *k*, the asymptotic behavior of the population can be described with the help of the branching process **Z**_*n*_ with offspring generating functions *G_i_* = *F_i+k_*, *i* ≥ 0, which only considers particles with type greater than *k*. To study this process we use results from the theory on decomposable branching processes (54), also used in (55) in the context of a viral population. The offspring mean matrix of the process is the upper triangular matrix 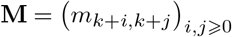, where *m_ij_* is the average number of *j*-mutants born from an *i*-mutant in the original process **X**_*n*_. Its largest eigenvalue is *m_k_* > 1, and **u** = (1, 0,…), **v** = ((*u/s_d_*)^*i*^/*i*!)_*i*≥0_ are respectively non-negative right and left eigenvectors of **M** corresponding to *m_k_*, satisfying **u · V**′ = 1. Assuming **Z**_0_ = (1,0,…), then 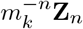 almost surely converges to *W*_**v**_, where *W* is a non-negative random variable with mean value 1, and {*W* = 0} = {*Z*_*n*,0_ → 0}. If **Z**_0_ = (*z*_0_, 0,…), then by independence of the lineages 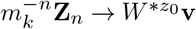, where *W*^\**z*0^ is the sum of *z*_0_ independent copies of *W*. The limit remains the same if **Z**_0_ = (*z*_0_, *z*_1_,…), since the *z_i_ i*-lineages with *i* > 0 have growth rate *m_k+i_* < *m_k_* and tend to 0 if normalized by 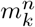. It follows that for any initial value **Z**_0_, 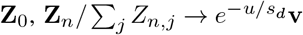 almost surely on the event 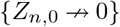.

Let 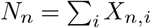 be the total population size at time *n* and let *f_i_* = *e^−u/s_d_^* (*u/s_d_*)^*i*^/*i*!. We deduce from what precedes that the fraction of *i*-mutants (*i* ≥ *k*) in the population satisfies

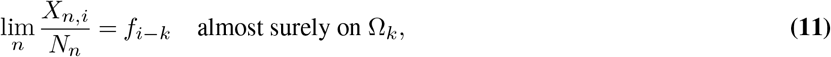

while this proportion converges to 0 if *i* < *k*. Since (Ω_*k*_)_0≤*k*≤*K*_ forms a partition of the survival set, the previous result can be stated as follows (setting the frequency to 0 if *N_n_* = 0):

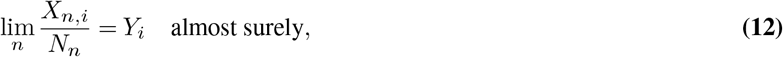

where *Y_i_* is a random variable equal to *f_i−k_* with probability *p_k_*(**n**), *k* ≤ min (*i, K*), and equal to 0 with probability 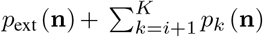. Therefore, the mean population fitness satisfies

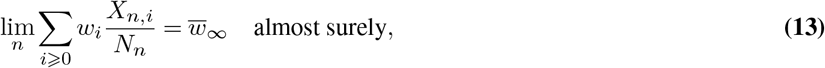

where 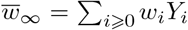 is a random variable equal to *w_k_e^−u^* with probability *p_k_*(**n**), 0 ≤ *k* ≤ *K*, and equal to 0 with probability *p*_ext_(**n**). It follows that the expected asymptotic population mean fitness is given by

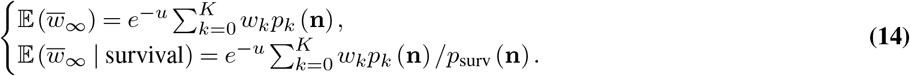

## Supplementary Note 2: Survival through multiple bottlenecks

### Markov chain model

In order to assess the effect of successive bottleneck events on the viral population, we introduce a stochastic model describing the evolution of the random composition of the population after each size reduction event. If the population reaches its carrying capacity *C*, a sample of size *B* is taken, resulting in a new founding population. As described earlier, the fate of this new population only depends on its initial numbers of supercritical mutants **n**, with 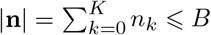 (the sample might include some particles which are not supercritical). We choose *C* large enough such that (*i*) the probability of not reaching *C* is close to the extinction probability of the population, (*ii*) if it reaches *C*, the mutant frequencies in the population follow the asymptotic random distribution Eq. (12). The evolution of the viral population going through these multiple size reductions is then entirely determined by the successive states **n**_0_, **n**_1_,…, where **n**_*b*_ describes the supercritical fitness classes after *b* bottleneck events. We describe this random evolution via a Markov chain with state space 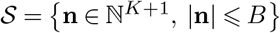, and transition probability from state **m** to state **n**:

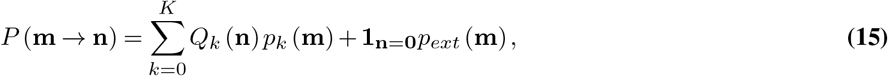

where *Q_k_*(**n**) is the probability of obtaining **n** supercritical mutants in the sample of size *B*, if the fittest surviving individuals in the population founded by **m** are of type *k* (event of probability *p_k_* (**m**)). It follows from Eq. (11) that on the latter event, the asymptotic proportion of non-supercritical mutants is 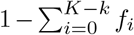, while the asymptotic proportion of supercritical *i*-mutants is *f_i−k_* if *i* ≥ *k*, and 0 otherwise. The random composition of the sample of size *B* thus follows a multinomial distribution with *B* trials and success probabilities 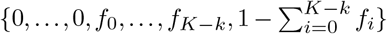, the last coordinate corresponding to the non-supercritical mutants. Therefore, the probability of getting **n** supercritical mutants is

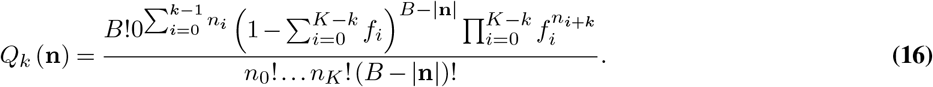

### Extinction probability under multiple bottlenecks

Let 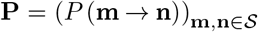 the stochastic matrix of this Markov chain, and **P**^*b*^ its *b*-th power. Then the probability for a viral population with initial state **n**_0_ to become extinct after going through at most *b* bottlenecks is

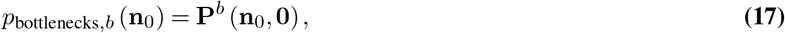

*i.e*. entry (**n**_0_, **0**) of matrix **P**^*b*^. This probability can thus be computed explicitly as soon as the extinction and click probabilities *p*_ext_ and *p_k_* are known, which is the case if for instance the numbers of offspring and of accumulated mutations follow Poisson distributions (see Eq. (24)).

## Supplementary Note 3: Epidemiology of genetic fragility

### Incubation period distribution

How long does it take for the in-host population carried by an exposed individual *E* to reach size *C*? Assuming *C* is large enough, this random time can be approximated thanks to the asymptotic behavior of the population. Let **n** the initial numbers of supercritical viral particles in the in-host population. As described earlier, if the population does not go extinct, its limit behavior depends on the type *k* of its fittest surviving particles. The distribution of the incubation period thus depends on **n** and *k*. When a host is infected, we therefore specify these parameters **n** and *k* and denote by *τ*_**n**,*k*_, *E*_**n**,*k*_ and *I*_**n**,*k*_ the corresponding incubation period, exposed and infectious states.

More specifically, when a susceptible *S* meets an infectious individual *I*_**m**,*l*_, it receives **n** supercritical viral particles with probability *Q_l_*(**n**), 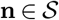, given by Eq. (16). The susceptible individual then either remains in state *S* with probability *p*_ext_(**n**), or enters in exposed state *E*_**n**,*k*_ with probability *p_k_*(**n**), 0 ≤ *k* ≤ *K*, given by Eq. (1). If so, it remains in *E*_**n**,*k*_ for *τ*_**n**,*k*_ time steps Eq. (5) before entering infectious state *I*_**n**,*k*_. Let *N_n_* the total viral population size carried by *E*_**n**,*k*_ at time *n*. We deduce from previously mentioned results that 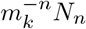 converges almost surely to *e^u/s_d_^ W*_**n**,*k*_, where *W*_**n**,*k*_ is a positive random variable whose distribution is given by Eq. (20), and explicitly by Eq. (26) in the particular case of Poisson distributed numbers of offspring and mutations described by Eq. (6). For *C* large enough, the time needed for *N_n_* to reach size *C* can consequently be approximated by

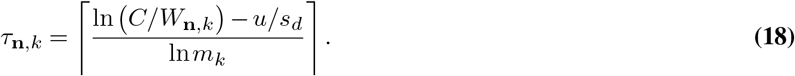

In the general setting, the random variable *W*_**n**,*k*_ is approximately the sum of *L* independent copies of *V*, where (i) *L* ≥ 1 is the random number of surviving *k*-lineages (*i.e*. a line of descent of a *k*-mutant with no *k*-mutants as ancestors), and (*ii*) *V* is the limiting distribution of 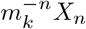 on its survival set, if (*X_n_*)_*n*_ is a monotype branching process modeling only *k*-mutants. Although not all explicit, we can state the following general results on *L* and *V*.

i. Let *k*_0_ the fittest initial type in **n**. If Muller’s ratchet does not click, *i.e*. if *k* = *k*_0_, then the number *L* of surviving *k*-lineages is at least 1 and at most *n_k_*. Since the survival probability of a *k*-lineage is 1 − *q_kk_*, *L* follows a binomial distribution with parameters *n_k_* and 1 − *q_kk_*, conditioned on being positive. If *k* > *k*_0_ however, there might be more than *n_k_* surviving *k*-lineages (namely also those stemming from the 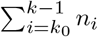 initial mutants). We therefore approximate the maximum number of surviving *k*-lineages by the average number of *k*-mutants born from **n** after one time-step, conditioned on the event that the surviving individuals are of type *k* (which we denote Ω_*k*_). This average is equal to 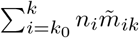, where 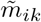 is the mean number of *k*-mutants born from one *i*-mutant, conditionally on Ω_*k*_. In the following computation, the subscript *i* indicates that the process starts with one *i*-mutant, and **q**_*k*_ stands for the infinite vector (*q*_*k*0_,…, *q_kk_*, 1,…) defined in Eq. (9). It comes from Eq. (7) and Eq. (1) that 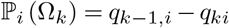, and that for any infinite vector **x**, using the notation 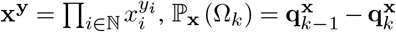. With this notation, 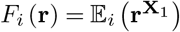 and 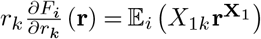. As a result, the mean 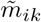 is obtained as

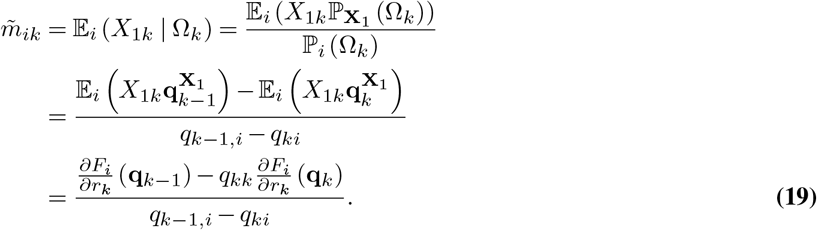
ii. Let (*X_n_*)_*n*_ a branching process with *X*_0_ = 1 and generating function *F*(*r*): = *F_k_*(*r*, 1, 1,…), *r* ∈ [0, 1]. It is therefore supercritical with mean offspring number *m_k_* > 1. From the theory on monotype branching processes (52) we know that its extinction probability is the smallest nonnegative solution of *F*(*r*) = *r*, which is by definition *q_kk_* given by Eq. (9). Moreover, 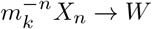 almost surely, where *W* is a nonnegative random variable whose Laplace transform 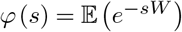 is the unique solution of *φ*(*m_k_s*) = *F*(*φ*(*s*)). Then the desired random variable *V* is distributed as *W* conditioned on *W* > 0. Since 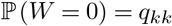, the Laplace transform of *V* is *φ*(*s*) = (*φ*(*s*) − *q_kk_*) / (1 − *q_kk_*).

To summarize, we assume that the random variable *W*_**n**,*k*_ appearing in Eq. (5) is given by

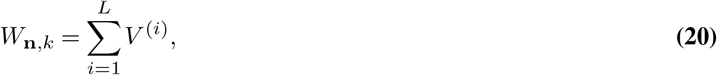

where *L* is a random integer such that

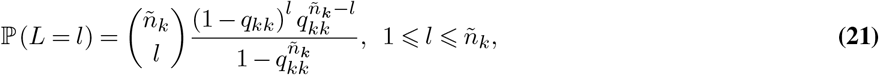

with

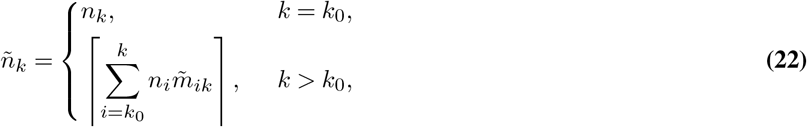

and where the *V*^(*i*)^ are independent copies of the random variable *V* with Laplace transform

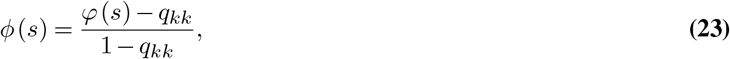

*φ* being the unique Laplace transform solution of *φ*(*m_k_s*) = *F_k_*(*φ*(*s*), 1,1,…)

## Supplementary Note 4: Explicit formulas

### Poisson distributed number of offspring

In the particular case of a Poisson distributed number of offspring and mutations (with generating functions Eq. (6)), an exact computation of (*q*_*k*0_,…, *q_kk_*) can be found, leading to explicit expressions of the extinction and Muller’s ratchet click probabilities Eq. (9)-Eq. (1) and of the expected asymptotic population mean fitness Eq. (2). Indeed, system Eq. (8) becomes

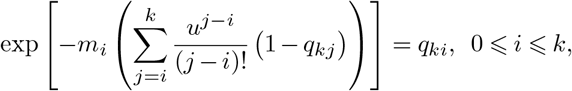

leading to

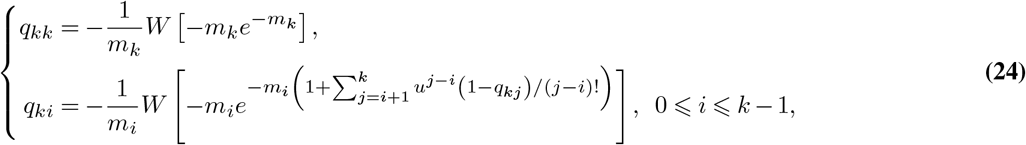

where *W*[·] stands for the principal solution of the Lambert *W* function.

In this particular case, we can also compute Eq. (19) and Eq. (23), and therefore provide the explicit distribution Eq. (20) of *W*_**n**,*k*_ involved in the incubation period of the exposed state Eq. (5). Indeed, with *F_i_* given by Eq. (6), we have 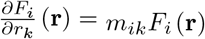, where *m_ik_* = *w_i_e^−u^u^k−i^*/(*k* − *i*)!. In addition, since **q**_*k*_ satisfies by definition *F_i_*(**q**_*k*_) = *q_ki_*, Eq. (19) becomes

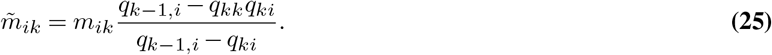

Finally, we show that, approximately, the random variable *V* involved in Eq. (20) follows an exponential distribution with parameter 1 − *q_kk_*. Indeed, the latter means that 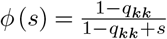, and thus 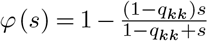. We have on the one hand

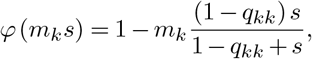

while on the other hand

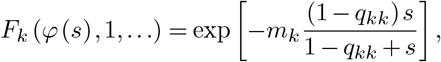

hence for *m_k_* close to 1, *φ*(*m_k_s*) ≈ *F_k_*(*φ*(*s*), 1,…), leading to our rough approximation. As the sum of independent exponentially distributed variables, *W*_**n**,*k*_ thus follows a Gamma distribution with shape and rate parameters

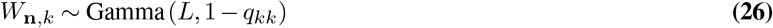

where *L* is given by Eq. (21) with

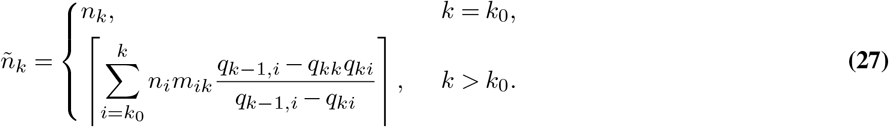

### Binary reproduction

Other reproduction mechanisms can be considered, with the same key assumption that the average number of (*i* + *j*)-mutants produced by an *i*-mutant is *w_i_e^−u^u^j^*/*j*!. For example, we can assume a binary reproduction (although less relevant for viral populations) where an *i*-mutant either dies, or produces two offspring with probability *w_i_*/2, each of them accumulating a random number of additional mutations following 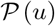. In this case, the generating function is quadratic

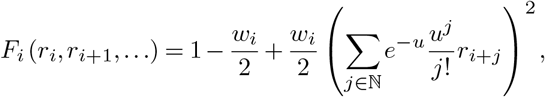

and system Eq. (8) comes down to

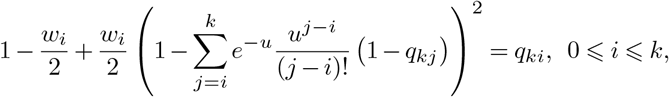

which can be solved explicitly as well.

